# Bacterial whole-cell biosensors illuminate spatially variable sialic acid availability within the inflamed mammalian gut

**DOI:** 10.1101/2024.10.23.619804

**Authors:** David Carreño, Clare M Robinson, Raphaella Jackson, Peirong Li, Vanessa Nunes, Susana A. Palma-Duran, Emma Nye, James I. Macrae, David T. Riglar

**Author notes:** Current Address: A*Star Genome Institute of Singapore, Singapore (Clare M Robinson). Current Address: Department of Women and Children’s Health, King’s College London, UK. Current Address: Department of Food Science, Research Centre in Food and Development A.C., Hermosillo, Mexico.

## Abstract

Host mucin-derived sialic acids are key drivers of microbial colonisation, growth and pathogenicity within the mammalian gut. However, their study is hindered by complex spatiotemporal dynamics: Many gut metabolites, including sialic acids, are rapidly consumed, transformed or absorbed by the microbiota or host, meaning that conventional measurements of faeces or bulk digesta often fail to capture their true local availability. In contrast, engineered bacterial-whole cell biosensors provide an *in situ* read-out of metabolite exposure at the point and time of use, before these molecules are depleted. Here, we demonstrate increases in the bioavailability of sialic acid in the inflamed mouse gut using an engineered *Escherichia coli* biosensor that reports on sialic acid exposure via NanR-regulated transcriptional circuit. The biosensor robustly colonises the mouse gut and remains functional for at least six weeks. Through organ-scale imaging at single bacterial resolution we observe strong correlations between disease status and biosensor response in two inflammation models. Profiling along the length of the gut uncovers regional variations between the maximal sialic acid sensing and the peak inflammatory response within host tissues in a murine colitis model. Simultaneous tracking of host and microbial markers of inflammation informs the therapeutic response to sialidase inhibition, which accelerates disease recovery. Together, these data illuminate the complex spatial dynamics involved in shared host-microbiome metabolism and demonstrate the broader power of using engineered bacterial biosensors to monitor *in situ* bioavailability of rapidly turned-over metabolites within the gut.

## Introduction

The metabolic and microbial composition of the mammalian gut is profoundly altered in disease conditions. Inflammatory gut environments have been broadly associated with increased prevalence of facultative anaerobes ^1^ and a decrease in microbiome-associated metabolite diversity ^2^. However, the site of sampling is critical for understanding. Quantification of various gut metabolites is complicated by their rapid depletion, reaction or reabsorption, altering local concentrations. Similarly, microbial disease signatures are often only evident in biopsy samples of the mucosal-associated microbiota ^3–5^. Thus, measurements from faecal sampling alone often provide a distorted view of the microbial and metabolic conditions that microbes experience *in situ*.

Mucus-derived carbohydrates are a particularly important nutrient class for microbes within the mammalian gut. The gut epithelium is protected by a layer of mucus, predominantly secreted by goblet cells. Highly glycosylated mucin proteins are a key component of this mucus, with MUC2 the dominant protein in both mouse and human ^6,7^. Variation in expression levels and the glycosylation patterns that decorate mucin proteins occurs throughout the gut, leading to structural and functional differences in mucus and altering its interactions with the gut microbiota ^8^. One key difference is the terminal monosaccharide of the glycosylation chain, which is typically either a sialic acid or fucose. Sialic acids are a family of nonulosonic acids that includes many derivatives of the basic neuraminic acid 9-carbon monosaccharide with N-acetylneuraminic acid (Neu5Ac) the most abundant. Data is limited but opposing gradients of sialylation and fucosylation are reported along the length of the colon in humans and mice ^9,10^.

Sialic acid monosaccharides can be liberated from mucin glycoproteins by microbial- and host-derived sialidase enzymes. While some bacterial species encode genes both for sialidases and sialic acid catabolism (eg. *Akkermansia muciniphila*), others typically express only sialidase (eg. *Bacteroides thetaiotaomicron* and *Phocaeicola vulgatus*) or catabolic gene repertoires (eg. *Escherichia coli* and various pathogenic species) ^11–14^. Thus, free sialic acid acts as a community resource that can be used by various species within the gut.

Aberrant sialylation, sialidase activity and serum sialic acid levels have been linked with various diseases including inflammatory conditions and cancer ^15,16^. Sialidase inhibition or knock-out of the α-2,3 sialyltransferase *St3gal4* reduces inflammation and perturbations to the microbiome otherwise caused by a dextran sulfate sodium (DSS) mouse colitis model ^17,18^. Huang and colleagues concluded that increased free sialic acid availability led to increased growth and persistence of *E. coli* bacteria within the gut, potentiating the downstream inflammatory phenotypes ^17^. However, direct quantification of sialic acid within both faeces and dissected gut contents have not shown increases in these models, likely due to rapid catabolism by the gut microbiota.

Sialic acid availability modulates bacterial gene expression, including induction of sialic acid catabolism genes, in various species. In *E. coli,* sialoregulon expression is controlled by the transcriptional regulator NanR ^19^. NanR represses gene expression of several operons, with direct binding by Neu5Ac leading to derepression and gene expression ^20^. In *Citrobacter rodentium,* sialic acid supports growth, directs chemotaxis, and induces virulence gene expression ^21^. On a biotechnology level, sialic acid was recently used to regulate a ‘sense-and-respond’ therapeutic engineered in *E. coli* bacteria through the *nanA* gene promoter ^22^. While the overall importance of sialic acid and sialidase activity is clear, there remains limited understanding of the spatial dynamics of this complex system.

Our group and others have demonstrated the power of engineered bacterial transcriptional reporters as robust, sensitive biosensors of the gut environment. Here, we combine the development of one such biosensor, induced by sialic acid, with organ-scale imaging to assess its activity within the gut. Assessing reporter activity in mouse DSS inflammation and *C. rodentium* infection models we show disease-specific biosensing which correlates microbial sialic acid sensing with host immune activity. By facilitating spatial analysis our sensor affords insights into regional segregation between host and microbial signals of inflammation. Finally, using *in situ* sensing we gain insights into the anti-inflammatory impacts of sialidase inhibition. The understanding gained illuminates *in situ* sialic acid levels as a microbial marker of inflammation and reveals previously unexplored spatial distinctions between microbiome and host inflammatory repertoires within the gut.

## Results

### Sialic acid impacts colitis development in a mouse DSS inflammation model

Microbiota-derived sialidase activity has been linked to alterations in both inflammation and microbiome composition, including in a dextran sodium sulphate (DSS)-induced inflammation model ^17,18^. To confirm these findings, we administered DSS (3%, 1%, or control) to C57BL/6 mice (n=3 per experimental group) in their drinking water for a period of 3 (3% group) or 7 (1% group) days (Figure 1A). To inhibit sialidase activity, a subset of animals additionally received the sialidase inhibitor Neu5Ac2en (IN) ^23^ via daily oral gavage (Figure 1A). Disease progression was monitored daily (Figure 1B-D). As expected, DSS provision led to increased inflammation as evidenced by elevated faecal LCN-2 and faecal occult blood levels (Figure 1C-D) and decreased colon length upon dissection (Figure 1E). Consistent with previous work, these impacts from DSS were significantly mitigated by sialidase inhibition, with inflammatory markers from both the 3% DSS + IN group at day 3 and 1% DSS + IN group at day 7 indistinct from those of controls (Figure 1C-E). Together these data confirm the importance of sialidase activity and sialic acid for inflammation in the DSS model.

**Figure 1.**
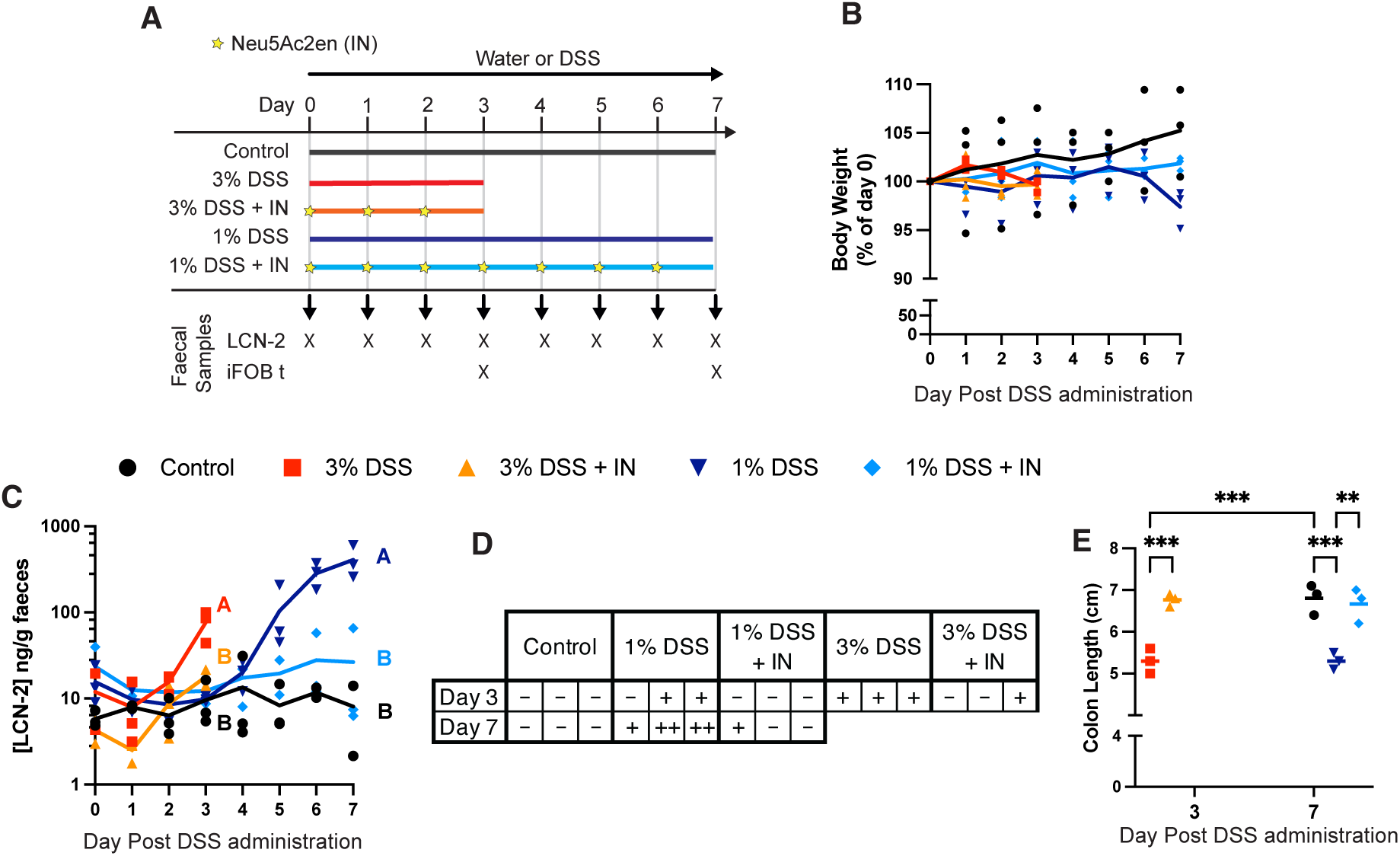
The impacts of sialidase inhibition in a DSS induced colitis mouse model. A) DSS at 3%, 1%, or 0% (control) was administered to C57BL/6 mice (n=3 per group) in the drinking water for a period of 3 (3% group) or 7 (1% group) days. To repress sialidase activity, some animals also received daily Neu5Ac2en sialidase inhibitor (IN) via oral gavage (stars). B) Body weight change, expressed as a percentage of starting body weight of each mouse. Line shows mean. C) LCN-2 levels as quantified by indirect ELISA of faecal pellets. Lines show mean. Statistics were calculated by one-way ANOVA with Tukey multiple comparison correction on AUC across all days to day 3 (3% DSS groups and control) or day 7 (1% DSS groups and control). Different compact letter (A-B) indicates significant differences between means. D) Faecal occult blood test (iFOB t) results from day 3 and 7. Absence (-) and presence of low (+) to high (++) levels of colorimetric change are indicated. E) Colon length as measured at the experimental endpoint on day 3 or 7. Line shows mean and statistics calculated by one-way ANOVA with Šídák multiple comparison correction. Adjusted p values are indicated *=p <0.05, **=p <0.005, ***=p <0.001.

The finding of delayed DSS-induced inflammation upon sialidase inhibition was further confirmed upon experimental repeat, this time delivering 3% DSS ± IN over 6 days (Supplementary Figure 1A-G). With the longer exposure to high level DSS in this experiment there were indications that inflammation was developing by days 5 and 6 even in the presence of the Neu5Ac2en inhibitor (Supplementary Figure 1B). Microbiome analysis of these animals by longread 16s rRNA sequencing at day 3 and day 6 also showed DSS-associated shifts driven by several bacterial species typically associated with inflammation, including elevation of key species expected to encode sialidases (Supplementary Figure 1D-G).

### Development of a live bacterial bioreporter to quantify sialic acid bioavailability to the microbiome

Despite its importance in inflammation, the dynamics of free sialic acid in the gut remain unclear. We reasoned that a microbially-encoded *in situ* biosensor could be a more reliable and accurate reporter of metabolite bioavailability within the gut microbiome given that rapid utilisation is expected to result in inaccurate quantification by traditional methods.

To develop a sialic acid-responsive whole cell biosensor, we cloned a small library of 8 putative sialic acid transcriptional reporters in the mouse commensal *E. coli* strain NGF-1 ^24^. The sensors tested included all *E. coli* promoter regions with predicted regulation by NanR ^20^: *nanATEKQ*, *nanCMS*, *nanXY* and *fimB*. Additionally, 4 synthetic promoters previously engineered with the NanR binding motif were tested ^25^. Sensors consisted of an inducible fluorescence output (mVenus) to assess sialic acid response combined with a constitutively expressed fluorophore (mKate2) for engineered bacterial identification (Figure 2A: Supplementary Figure 2A). All circuits were integrated at single copy in the genome to preserve native NanR sensitivities since previous studies have noted loss of NanR regulation when using plasmid-based reporter systems ^22^. Despite lowering biosensor output, this design choice should also minimise impacts of the engineered circuit on the bacterial chassis and thus facilitate robust growth and function within the mouse gut.

**Figure 2.**
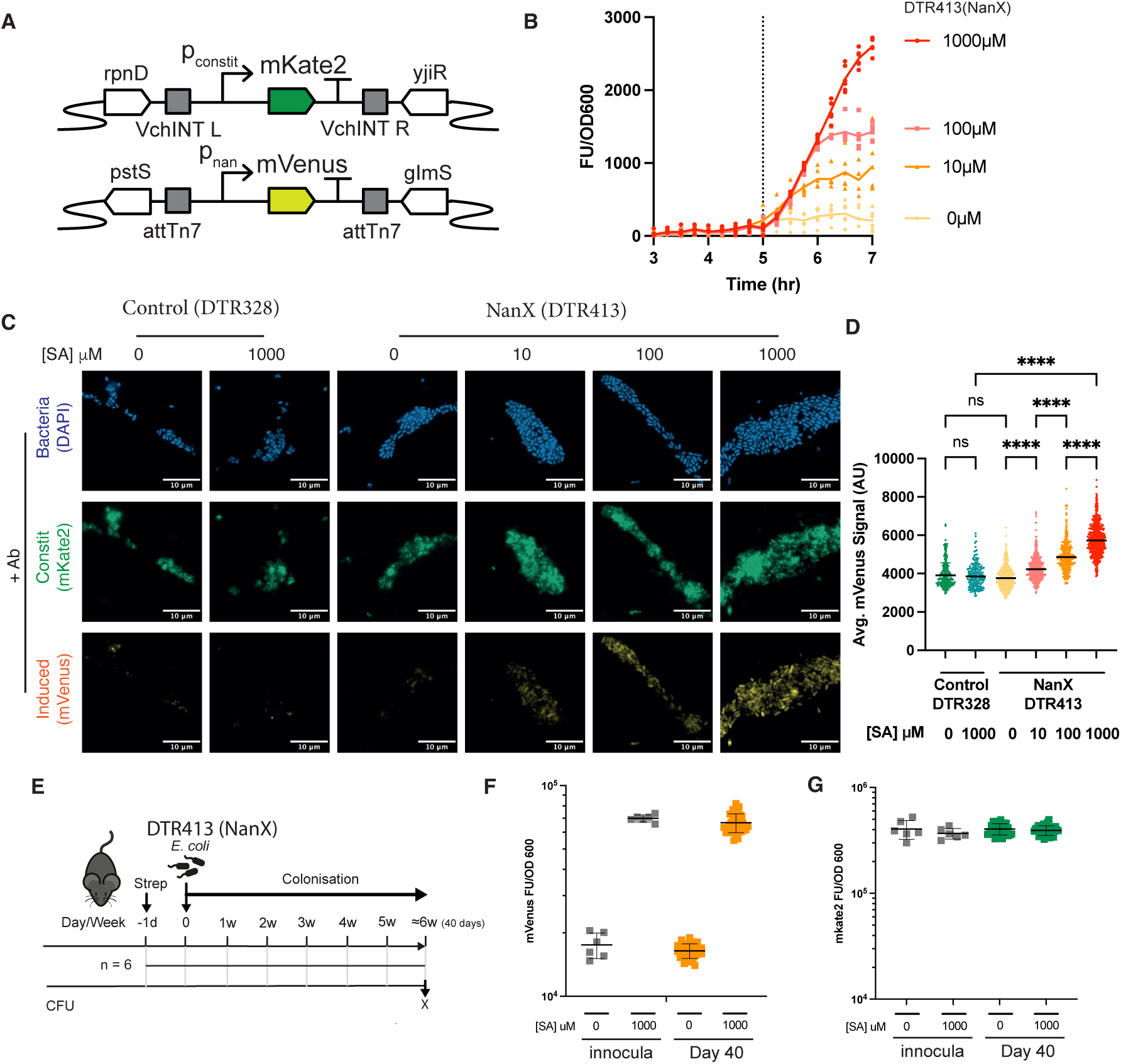
Development of a live bacterial whole-cell biosensor shows quantitative response to sialic acid bioavailability. A) Gene circuit construction of fluorescent sensor strain. *E. coli* NGF-1 bacteria were engineered with a constitutively expressed mKate2 fluorescent reporter, inserted into the genome between *rpnD* and *yjiR* genes. Inducible promotors predicted to be controlled by *E. coli NanR* gene regulator were cloned in front of an mVenus fluorescent reporter and inserted into the genome downstream of the *glmS* gene. B) Fluorescence output of DTR413 (NanX), normalised by OD600 and DTR328 (negative control) background, following mid-growth Neu5ac induction (0μM, 10μM, 100μM or 1000μM) in M9 media + 0.4% glycerol. Graph shows mean of 6 replicates. The dotted line represents the time of Neu5Ac addition. C) Widefield microscopy-based detection of antibody-labelled fluorescent reporter expression from DTR413 (NanX) and DTR328 (negative control) in response to the induction with increasing concentrations of Neu5Ac for 6h in LB medium. D) Average mVenus signal (AU) of segmented bacteria from C. Statistics shown for Kruskal Wallis test with Dunn’s multiple comparison adjustment as measured between treatments shown. * = p<0.05, ****= p<0.0001. E) DTR413 sialic acid-inducible whole cell biosensors were administered to C57BL/6 mice (n = 6) by gavage 24 hours following a single dose of streptomycin (5mg per mouse by oral gavage). After 40 days of gut colonisation, 36 colonies from 6 animals were isolated from faecal pellets to perform an *ex vivo* analysis. F) Fluorescence mVenus output of individual colonies exposed to 1000 uM of Neu5Ac and normalised by OD600 at 6 hours. G) Fluorescence mkate2 output of individual colonies, performed in parallel with F. Graph shows mean of 6 replicates.

Growth of each biosensor in the presence or absence of Neu5Ac (SA; 0μM, 10μM, 100μM or 1000μM) identified two sensors with increased fluorescence response at higher sialic acid levels (Supplementary Figure 2B). In particular, the *E. coli nanXY* gene promoter (P*_nanX_*) (strain: DTR413) showed the desired characteristics of low background output and the highest proportional activation so was chosen for all further testing. Induction of DTR413 at mid-log phase demonstrated the strain’s rapid response dynamics, with increased fluorescence detected as early as ∼30 minutes post induction and distinction between all sialic acid concentrations tested by ∼1-1.5 hours (Figure 2B). Mirroring the overall specificity and selectivity of *E. coli* for different sialic acid variants, DTR413 showed similar sensitivity to Neu5Ac and 5-glycolylneuraminic acid (Neu5Gc), and minimal sensitivity to 2-keto-3-deoxy-nononic acid (KDN) (Supplementary Figure 2C). By comparison, reporter response was not impacted by N-acetylmannosamine (manNAc) (Supplementary Figure 2C) or the Neu5Ac2en sialidase inhibitor (Supplementary Figure 2D).

### Microscopy-based detection of biosensor response

We next confirmed the ability to detect DTR413 reporter activation by microscopy. In preparation for imaging of the gut, where the combination of inconsistent oxygen levels and a highly auto-fluorescent background create challenges for direct fluorescence detection, we amplified reporter signals through indirect immunohistochemistry (IHC). DTR413 bacteria were induced with Neu5Ac for 6 hours, then fixed, permeabilised and labelled with *α*-GFP (mVenus), *α*-RFP (mKate2) and DAPI prior to widefield imaging. Reporter induction was readily detectable and exhibited a clear increase with escalating sialic acid concentrations (Figure 2C). To quantify the induction, image analysis was performed on bacteria segmented using a classifier generated using Labkit software in ImageJ ^26^. Immunohistochemistry amplified bacteria showed mean induction of mVenus proportional to Neu5Ac (Figure 2D), but no systematic impact of Neu5Ac on mKate2 levels (Supplementary Figure 2E). Together these data demonstrate the ability for this engineered biosensor to report on sialic acid availability.

### Sialic acid biosensor function during colonisation of the murine gut

To assess the long-term functionality of the sialic acid biosensor in vivo, we next designed an experiment in which mice were colonised with DTR413 following a single low dose of streptomycin and maintained for 40 days before analysis (Figure 2E-G). We have previously shown that engineered *E. coli* NGF-1 can colonise the murine gut at 10^4–6^ CFU/g faeces for extended periods after a single 5 mg streptomycin dose ^27,28^. DTR413 established stable long-term colonisation and maintained unchanged reporter function and sensitivity over 40 days in the gut based on *ex vivo* analysis of colonies (36 colonies from 6 animals). Together this indicates high genetic and functional stability of the genomically integrated biosensor (Figure 2F-G).

### Spatial profiling indicates regional variations in sialic acid availability within the gut

To assess biosensor response to inflammatory conditions, we delivered DTR413 bacteria to mice (n = 4 per group) as previously, with or without 3% DSS in drinking water and daily Neu5Ac2en sialic acid inhibitor gavages (Figure 3A). Markers of disease progression mirrored those in previous experiments (Figure 3B-J, Supplementary Figure 3D-H), again demonstrating reduced inflammation in animals co-administered with the Neu5Ac2en sialidase inhibitor (Figure 3D).

**Figure 3.**
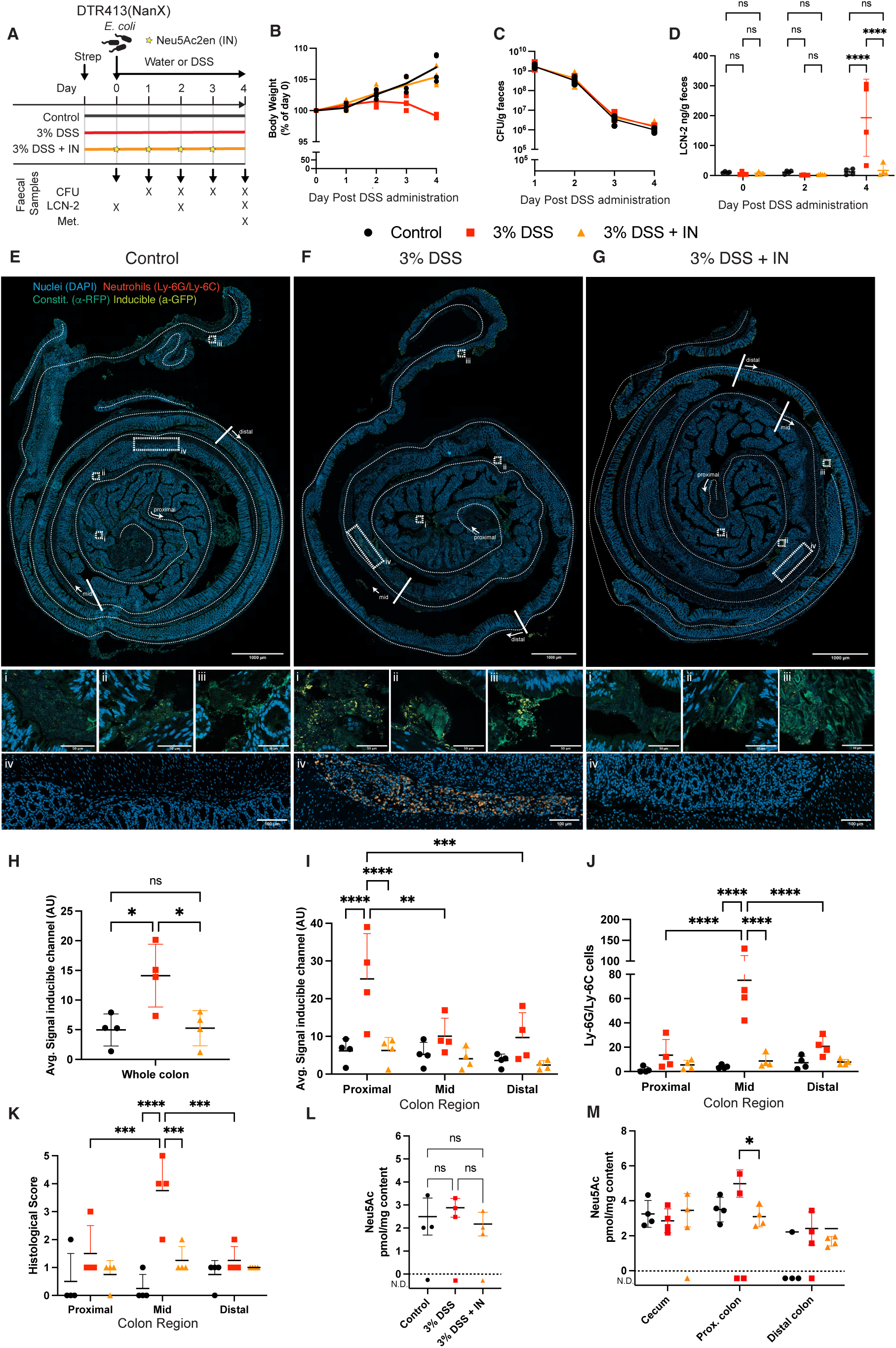
Longitudinal spatial availability of sialic acid during inflammation in a colitis mouse model. A) DTR413 sialic acid-inducible whole cell biosensors were administered to C57BL/6 mice (n = 4 per group) by gavage 24 hours following a single dose of streptomycin (5mg per mouse by oral gavage). DSS was administered from the day of bacterial delivery in the drinking water at 0% (control) and 3% with or without daily oral gavage of Neu5Ac2en sialidase inhibitor (IN; stars). B) Body weight change, expressed as a percentage of starting body weight of each mouse. Line shows mean. C) DTR413 abundance was measured in faecal pellets collected daily post administration. D) LCN-2 levels, as quantified by indirect ELISA in faecal pellets collected at day 0, 2 and 4. Graph shows mean ± SD. Statistics show results from two-way ANOVA with Tukey multiple comparison correction between samples on each day as shown. ****=p<0.0001. E-G) Representative tile images of whole colon from E) Control, F) 3% DSS, and G) 3% DSS + IN treated animals. IHC labelling of inducible mVenus (α-GFP; yellow), constitutive mKate2 (α-RFP, green), and neutrophils (Ly-6G/Ly-6C, orange) along with nuclear staining (DAPI, blue) are shown. Annotations indicate the divisions used for longitudinal sections of proximal, mid and distal colon. Representative zooms of proximal (i), mid (ii) and distal colon (iii) for reporter-gene labelling, and mid colon for neutrophil infiltration detail (iv) are shown also. Scale bars are shown in figure. H) Geometric means of average signal of inducible mVenus reporter intensity (α-GFP) within bacteria segmented for constitutive (α-RFP) fluorochrome on the whole colon swiss-roll at day 4. I) Geometric mean of inducible mVenus reporter intensity (α-GFP) for the longitudinal spatial profile across the colon, with the swiss-roll subdivided into proximal, mid, and distal regions for regional comparison at day 4. J) Neutrophil counts in gut regions based on α-Ly-6G/Ly-6C immunolabelling. K) Histological Score in each gut regions from blinded analysis of H&E labelled sections, combining a sum of independently scored changes in crypt architecture, inflammatory infiltrate, muscle thickening, goblet cell depletion and crypt abscess. L) Concentration of Neu5Ac in faecal pellets at day 4 as analysed by GC-MS. M) Neu5Ac concentration in caecum, proximal and distal colon at day 4 as analysed by GC-MS. Graphs show mean ± SD from 4 replicate animals. Statistics show results from one-way ANOVA (H,L) or two-way ANOVA (D, I-K, M) tests with Tukey multiple comparison adjustment comparing experimental groups within a gut region and between gut regions within each experimental group. * = p<0.05, **=p<0.005; ***p<0.0005, ****p<0.0001. For L-M, not determined (ND) values represent GC-MS chromatograms that failed quality control assessment and were excluded from further calculations; they are shown here for completeness only.

For biosensor analysis, dissected gut samples were prepared as ‘swiss-rolls’ (Supplementary Figure 3A). Sectioned samples were stained for constitutive (*α*-RFP to recognise mKate2) and sialic acid inducible (*α*-GFP to recognise mVenus) reporters, along with DNA (DAPI) and neutrophils (*α* Ly-6G/Ly-6C) and whole colons were imaged via confocal microscopy (Figure 3E-G). Inducible reporter levels in bacteria segmented based on constitutive reporter expression, demonstrated an inflammation-specific increase in sialic acid sensing (Figure 3H).

Given the spatially diverse nature of the gut environment, we further analysed these images in three equal sub-sections: proximal, mid, and distal colon (Figure 3I-K). Notably, reporter response in the proximal colon was significantly higher than in the mid and distal colon in these animals (Figure 3I). By comparison, several key indicators of host inflammation were concentrated in the mid colon region. Counts of neutrophil infiltration from the same histological sections were consistently elevated in DSS treated animals within the mid-colon region, with limited indication of polymorphonuclear cell clusters in proximal and distal colon of DSS treated animals, nor at any sites in control or DSS + sialidase inhibitor treated groups (Figure 3E-Giv; Figure 3J). Similarly, monocyte-derived cells (CD68) and leukocytes (CD45) including T cells (CD3), showed signs of elevation within the mid colon above controls and sialidase inhibitor treated animals (Supplementary Figure 3B-C). These results were mirrored by blinded disease severity scoring of hematoxylin and eosin (HE) stained adjacent sections (Figure 3K; Supplementary Figure 3D-F).

This finding of spatial distinction between host and microbial markers of inflammation was confirmed upon experimental repeat (Supplementary Figure 4). Importantly, this repeat was performed 6-weeks following initial bacterial delivery, demonstrating both the suitability of our sensing approach for reporting on steady-state conditions at natural commensal colonisation levels and ruling out any influence of antibiotic treatment on these findings.

Given the regional specificity of sialic acid biosensor response, we next asked whether conventional metabolite measurements could detect similar regional differences in sialic acid levels. Neu5Ac was therefore quantified in faecal samples and digesta along the dissected gut using gas chromatography-mass spectrometry (GC-MS). Free Neu5Ac levels detected in faecal samples were indistinguishable between treatment groups (Figure 3L). Analysis of regional Neu5Ac levels was complicated by variability in the amount and consistency of digesta between animals and treatment groups. Nevertheless, when digesta samples were extracted from broad regions of the gut (cecum, proximal and distal halves of the colon), elevated Neu5Ac levels were detected in the proximal colon of DSS treated animals, consistent with our whole-mount imaging results (Figure 3M).

Together our results point to increased sialic acid bioavailability during DSS-induced inflammation, but spatial separation between sites of sialic acid elevation in the gut lumen and the primary site of inflammatory injury within the gut. They also highlight the superiority of an engineered whole-cell biosensor over direct GC-MS based measurements for resolving regional differences in a rapidly turned-over metabolite.

### In situ sialic acid is a marker of inflammation across models of disease

We next asked whether sialic acid biosensing could similarly inform on disease burden in a distinct inflammatory setting. Like DSS, detection of free sialic acid levels in faeces during *C. rodentium* infection, a model for human pathogens enteropathogenic *E. coli* (EPEC) and enterohaemorrhagic *E. coli* (EHEC), is inconsistent and microbiome dependent ^21,29^.

To track *in situ* sialic acid bioavailability during *C. rodentium* infection we therefore delivered DTR413 bacteria to C57Bl/6 mice by oral gavage 24-hours following streptomycin treatment (5mg/mouse) (Figure 4A). Four days later, mice (n=6) were infected with *C. rodentium* ICC169. Over the subsequent 6 days of infection, body weight (Figure 4B) and faecal *E. coli* DTR413 bacteria (Figure 4C) remained consistent between infected and control (n=4) groups. By comparison, faecal *C. rodentium* ICC169 levels initially dipped, consistent with the well characterised establishment phase of infection when faecal CFUs routinely drop by up to ∼10^6^ ^30,31^, before elevating in the second phase of infection (Figure 4D). Citrobacter burden was also evident by imaging of dissected gut using a polyclonal α-CR antisera (Figure 4E). By day 6 post infection, infected animals also had a significant but modest elevation in faecal LCN-2 levels (Figure 4F).

**Figure 4.**
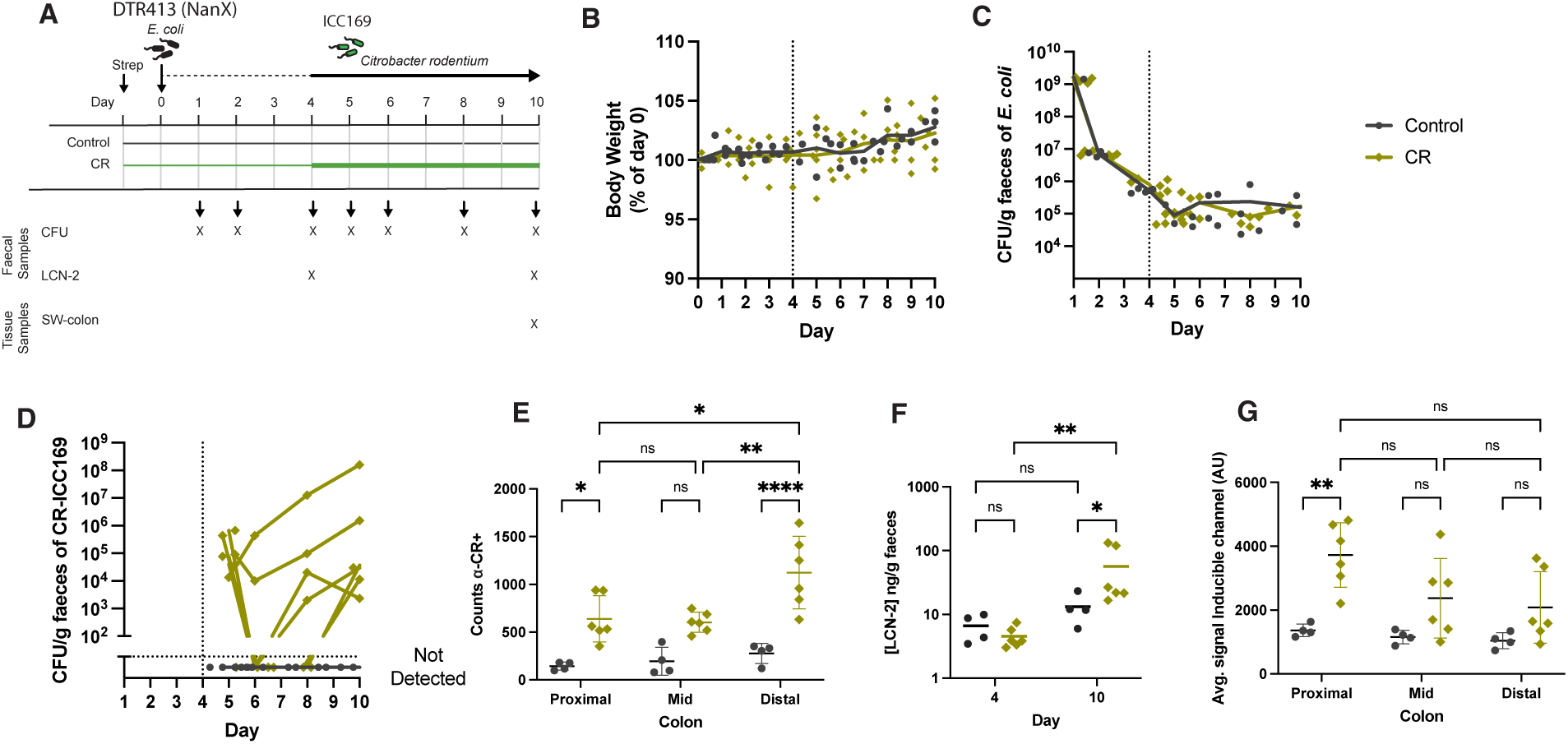
*Citrobacter rodentium* drives changes in bioavailability of sialic acid. A) DTR413 sialic acid-inducible whole cell biosensors were administered to C57BL/6 mice (n = 10) by gavage 24 hours following a single dose of streptomycin (5mg per mouse by oral gavage). After four days, 6 mice were challenged with 10^9^ CFU of *C. rodentium* ICC169 by oral gavage. Infection course lasted for six days. B) Body weight change, expressed as a percentage of starting body weight of each mouse. Line shows mean. C) DTR413 abundance was measured in faecal pellets collected at day 1, 2, 4, 5, 6, 8 and 10 post administration. D) ICC169 abundance was measured in faecal pellets collected at day 5, 6, 8 and 10 post administration of DTR413. E) *C. rodentium* (CR) counts in gut regions based on polyclonal anti-CR. F) LCN-2 levels, as quantified by indirect ELISA in faecal pellets collected at day 4 (before CR challenge) and day 10. G) Geometric means of average signal of inducible mVenus reporter intensity (α-GFP) within bacteria segmented for constitutive (α-RFP) fluorochrome on swiss-roll sections of proximal, mid and distal colon at day 10. For E-G and L, Line shows mean and statistics calculated by two-way ANOVA with Tukey’s multiple comparison correction. Adjusted p values are indicated *=p <0.05, **=p <0.005, ***=p <0.001.

Imaging of induced mVenus levels in DTR413 bacteria demonstrated elevated sialic acid sensing in infected animals (Figure 4G). Interestingly, while sialic acid response was again highest in proximal colon, unlike in the DSS model several individual animals also showed signs of high response in the mid- and distal-colon (Figure 4G). These findings demonstrate the utility of *in situ* sialic acid sensing as a measure of disease across different inflammatory models.

### In situ sialic acid biosensing tracks in vivo efficacy of sialidase inhibition

Finally, having established DTR413’s ability to report on *in situ* levels of sialic acid, we used this capacity to further explore the therapeutic impact of the Neu5Ac2en sialidase inhibitor. Previously, we and others have demonstrated the delay of inflammatory onset by Nue5Ac2en (Figure 1) ^17^. To now test the impact of Neu5Ac2en on acute inflammation, we delivered DTR413 to C57Bl/6 mice (n=5 per group) and compared between groups that had been treated with DSS for 5 days then allowed to recover for a further 5 days in the presence or absence of daily Neu5Ac2en sialidase inhibitor gavages (Figure 5A-B). Healthy and acute DSS (3% DSS for 5 days) treated groups were also included as controls (Figure 5A-B).

**Figure 5.**
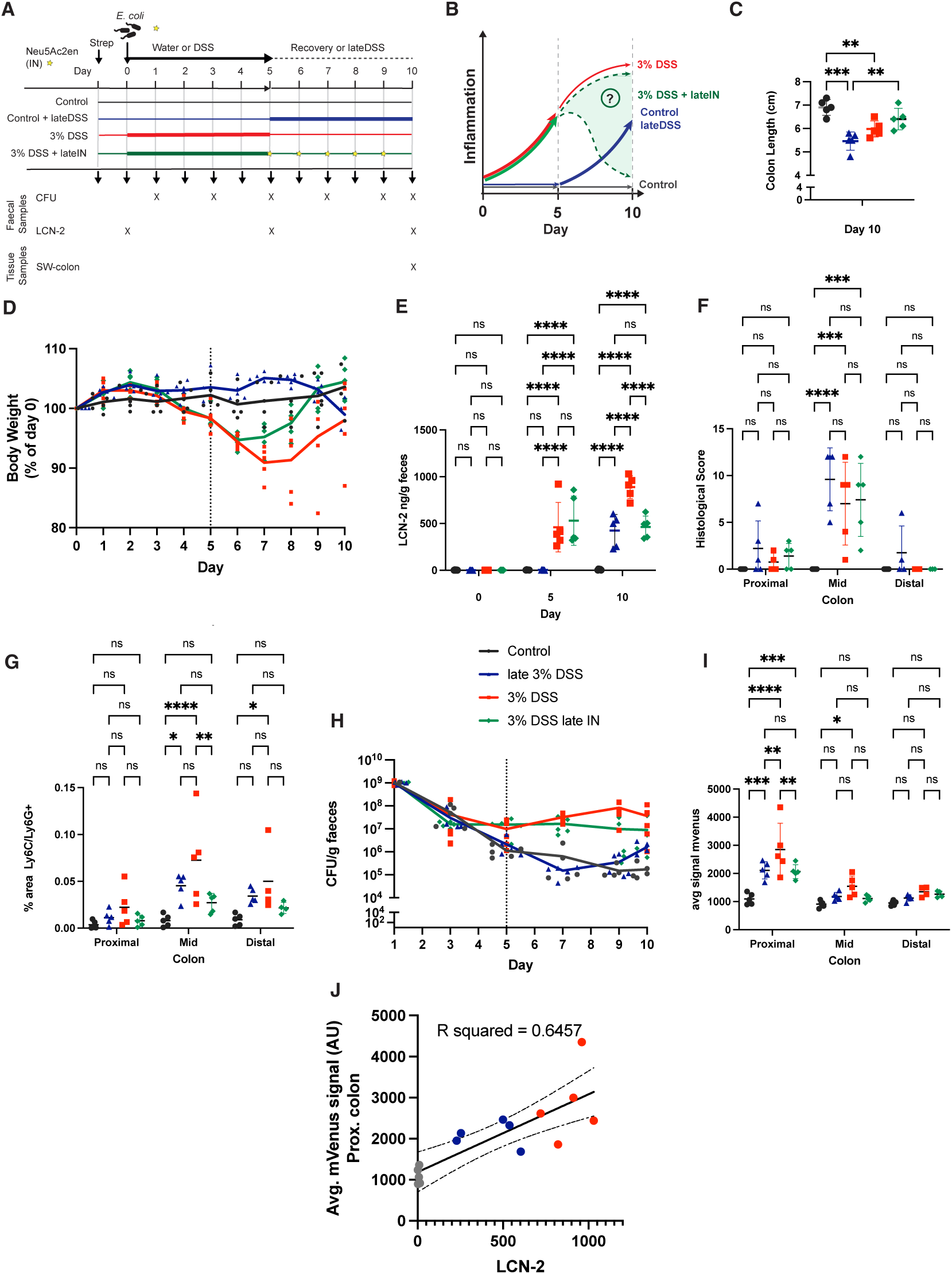
Sialic acid biosensing reflects disease burden across inflammatory states and identifies incomplete sialidase inhibition *in vivo*. A) DTR413 sialic acid-inducible whole cell biosensors were administered to C57BL/6 mice (n = 5 per group) by gavage 24 hours following a single dose of streptomycin (5mg per mouse by oral gavage). DSS was administered in the drinking water at 0% (control) and 3% for the last five days of experiment (late 3% DSS) or from the day of bacterial delivery with a daily gavage of sialidase inhibitor Neu5Ac2en (3% DSS+ late IN) or without (3% DSS). B) Expected inflammatory trajectory across the experimental timeline shown in A. The shaded region indicates the expected inflammation window targeted by the 3% DSS + inhibitor treatment course. C) Body weight change, expressed as a percentage of starting body weight of each mouse. Line shows mean. D) Colon length as measured at the experimental endpoint on day 10. Line shows mean and statistics calculated by one-way ANOVA with Tukey’s multiple comparison correction. Adjusted p values are indicated *=p <0.05, **=p <0.005, ***=p <0.001, ****p<0.0001). E) LCN-2 levels, as quantified by indirect ELISA in faecal pellets collected at day 0, 5 and 10. Graph shows mean ± SD. F) Histological Score in each gut regions from blinded analysis of H&E labelled sections, combining a sum of independently scored changes in crypt architecture, inflammatory infiltrate, muscle thickening, goblet cell depletion and crypt abscess. Graphs show mean ± SD. G) Percentage of area positive for neutrophils based on α-Ly-6G/Ly-6C immunolabelling in each of the colon regions. H) DTR413 abundance was measured in faecal pellets collected at day 1, 3, 5, 7, 9 and 10 post administration. I) Geometric means of average signal of inducible mVenus reporter intensity (α-GFP) within bacteria segmented for constitutive (α-RFP) fluorochrome on swiss-roll sections of proximal, mid and distal colon at day 10. For E-G and I, Line shows mean and statistics calculated by two-way ANOVA with Tukey’s multiple comparison correction. Adjusted p values are indicated *=p <0.05, **=p <0.005, ***=p <0.001, ****p<0.0001. J) Pearson’s correlation analysis by between avg. mVenus signal of the inducible fluorescence channel in the proximal colon and LCN-2 levels in faecal pellets (*r*=0.8036*, R*^2^*=*0.6457). Data from DSS +IN treated animals was not used in correlation analysis to avoid possible impacts from sialidase inhibition on results.

Animals receiving sialidase inhibitor showed broad signs of accelerated recovery from inflammation at day 10 compared with animals recovering in the absence of inhibitor based on colon length (Figure 5C), bodyweight (Figure 5D) and faecal LCN-2 (Figure 5E). While blinded scoring by a pathologist showed that histological signs of inflammation remained (Figure 5F), there was nevertheless evidence of tissue recovery noted in two inhibitor treated animals (Supplementary Figure 5) and α-Ly-6G/Ly-6C immunolabelling demonstrated reduced neutrophil infiltration in the mid-colon of inhibitor treated animals (Figure 5G). This indicates that sialidase inhibition can not only mitigate onset of inflammation but can also accelerate recovery from active DSS induced damage.

Analysis of DTR413 biosensor bacteria facilitated further interrogation of sialic acid bioavailability in each treatment group. As previously described, extended DSS regimens led to elevated DTR413 levels in the gut compared with untreated controls (Figure 5H). Again, sialic acid induced mVenus levels were particularly elevated with the proximal colon of animals (Figure 5I), with a strong correlation between host inflammation levels as measured by faecal LCN-2 and proximal sialic acid reporting across control and DSS treated animals (p=0.0003, *r =* 0.8036, *R*^2^=0.6457) (Figure 5J). Notably, recovery from DSS administration in the presence of Neu5Ac2en reduced the average mVenus signal compared with the DSS recovery alone group, in line with other inflammatory markers (Figure 5I). However, sialic acid sensing remained higher than baseline in Neu5Ac2en treated animals despite daily administration of the sialidase inhibitor, suggesting incomplete sialidase inhibition.

Together, these experiments show that sialidase inhibition speeds up recovery from acute inflammation, and that sialic acid availability within the proximal gut lumen is a strong correlate of host immune activity.

## Discussion

Here we have combined engineered live microbial whole-cell biosensors with imaging to provide *in situ* spatial understanding of sialic acid bioavailability within the gut. Despite the established importance of this pathway for host-microbiome interactions and pathogenesis of various diseases, direct detection of free sialic acid in the gut has always been complicated by its rapid utilisation by the microbiota. By comparison to faecal quantification, our biosensor’s *in situ* response to sialic acid was strongly correlated with several measures of inflammatory disease and host immune activation and clearly separated between treatment groups in both DSS colitis and *C. rodentium* infection models (Figure 3-4). Biosensing was sensitive enough to detect the low inflammation levels encountered relatively early in the onset of both models. Thus, our data further support the potential to use free sialic acid as a biomarker for inflammatory disease status in the gut, but only when it is sensed *in situ*.

Spatial analyses in the DSS model suggested regional differences between sialic acid availability, which peaked within the proximal colon, and the primary site of inflammatory impact within the host, the mid colon (Figure 3, 5). This finding was observed across all experiments in acute and recovery DSS treatment regimens (Figure 3-5) and in animals colonised with DTR413 6 weeks prior to testing, demonstrating that antibiotic treatment is not driving these results (Supplementary Figure 4). The regional importance of the mid-colon in the murine response to DSS has been described previously via spatial transcriptomics ^32^. Our findings point to a system-wide mechanism of host-microbiome interaction that drives availability of a key inflammation-linked nutrient for microbes at distinct sites to inflammatory tissue damage. Interestingly, while sialic acid response again correlated with disease burden during *C. rodentium* infection, the response was more spatially heterogeneous throughout the colon in several animals (Figure 4). Thus, while there are previously described gradients of mucus sialylation in the mouse gut with higher levels expected in proximal vs distal regions, this is insufficient to explain our findings.

The finding that daily administration of the Neu5Ac2en sialidase inhibitor accelerates recovery from active colitis is of interest in the context of anti-inflammatory therapies (Figure 5). Importantly, however, DTR413 sensing indicated that considerable sialic acid remained available within the gut of animals treated with sialidase inhibitor. Together with the eventual development of inflammation after 5-6 days of concurrent DSS and Neu5Ac2en delivery (Supplementary Figure 1), these data suggest that sialidase inhibition remains incomplete within these animals. One possible reason for this is the likely presence of sialidase enzyme subsets that are poorly inhibited by Neu5Ac2en, for example Bacteroides sialate O-esterases (EstA) ^33^. Similarly, inhibition of host sialidases by Neu5Ac2en is relatively weak, and Neu3 has been implicated in sialic acid removal from intestinal alkaline phosphatase ^34^. Alternatively, breakthrough sialidase activity could be a result of an insufficient dosing amount or frequency. The latter may explain the lack of impact seen in another study when inhibitors, including Neu5Ac2en were dosed less often ^33^. This points to potential strategies for optimising activity, with *in situ* sialic acid detection a powerful tool in this process.

These findings also have important implications for the emerging field of engineered live bacterial diagnostics and therapeutics, particularly those that aim to induce therapeutic response specifically at the site of disease ^35,36^. Indeed, sialic acid inducible circuits have been developed as prototype ‘sense-and-response’ systems for *in situ* delivery of therapeutics in antibiotic-induced inflammation^22^. Our results support the use of sialic-acid driven circuits to drive inflammation-specific functions, while at the same time highlighting the pitfalls of regional specific gene regulation of this circuit, which may be incompatible with many applications.

A particular design focus was the development of low-burden circuits with minimised reporter expression to achieve robust functionality when colonising the gut. The engineered biosensors developed here achieved effective colonisation similar to previous studies using the same *E. coli* host ^27,28^. DTR413 bacteria remained functional while growing unselected in the conventional mouse gut for over 40 days and were able to reliably report on *in situ* sialic acid levels throughout this period (Supplementary Figure 4). This is a fundamentally different approach to whole cell biosensing than using the typical high-expressing, plasmid-based reporters dosed for immediate response, which are typically lost or mutated over short periods in the body. One inherent challenge arising from this approach, however, is the lower signal output from integrated circuits. Future circuit optimisations, achievable through promoter engineering, synthetic circuit amplification, adaptive laboratory evolution, or equivalent approaches are promising avenues for amplifying reporter output while maintaining a genome integrated design that limits growth burden.

The imaging of microbial transcriptional reporters is a generalisable approach to dissecting metabolic pathways involving host and microbiota. Given the microbiome’s extensive sensing capacity, bacterial biosensors have been developed to detect a range of metabolites, and ongoing work in our laboratory and others aims to expand this repertoire ^28,37^. Direct metabolite detection, especially in faeces, may be unrepresentative when metabolites are also utilised by host and/or microbiota. There are also practical limitations in sample collection from localised regions of the gut, with amount and consistency of digesta varying widely between individuals and with disease. Indeed, this is the most likely cause of ‘not determined’ values in several of our regional colon GC-MS samples (Figure 4). This fundamentally increases variability and limits utility of results. Conversely, biosensing in the context of a live member of the microbiota can offer insight into the dynamics of availability for utilisation driven by the complex interplay of localised metabolite production and utilisation.

These tools promise to broaden the utility and understanding of key metabolites as biomarkers for health and disease with potential benefits both in research and the clinic.

## Materials and Methods

### Bacterial strain construction

Details of all bacterial strains developed for this study are provided (Reagents and Resources Table). All synthetic circuits were integrated into the chromosome of the mouse commensal *E. coli* NGF-1 strain carrying a rpsL lys42arg mutation which affords high-level streptomycin resistance^24,38^.

Constitutive fluorescence was achieved using an mKate2 variant, which includes 11 N-terminal residues from mCherry for increased translation ^39^, expressed downstream of the strong constitutive promoter pRNA-1. For integration in the genome, the pSL1521 pSPIN plasmid carrying the machinery for INTEGRATE mediated gene insertion ^40^ was modified to target the region between *rpnD (yjiP)* and *yjiR*. The choice of site was based on previous identification of this site as supporting high expression of an mCherry cassette without impacting *E. coli* growth ^41^. The crRNA was designed to target a PAM site conserved between *E. coli* NGF-1, MG1655 and Nissle 1917 strains. The crRNA sequence was ordered as single stranded oligonucleotides (IDT) and duplexed by resuspending oligos at 200 μM in annealing buffer (100 mM potassium acetate; 30 mM HEPES, pH 7.5; IDT), mixing forward and reverse oligos in equal volumes and incubating at 95° C for 2 mins followed by gradual cooling to 25°C over 45 minutes. The duplexed crRNA was then assembled into the pSL1521 vector by BsaI golden-gate assembly (GGA). pRNA1-mKate2 was amplified from LPT36 via PCR and J23101-mKate2 was ordered as a gBlock Gene Fragment (IDT), with both being assembled into the *rpnD-yjiR* targeted vector via HiFi assembly using manufacturer guidelines (NEB). Plasmids were first assembled in *E. coli* Dh5a, followed by miniprep and electroporation into *E. coli* NGF-1. Cells harbouring plasmids were grown overnight and colonies screened by PCR for genomic integration of fluorescent cassettes.

For rapid cloning of inducible mVenus constructs, promoters were assembled via SapI mediated GGA into pCR43, a modified Tn7 insertion vector derived from pGRG36 ^42^ via pCR06 ^28^ to facilitate GGA cloning of promoters to drive mVenus expression. Native *E. coli* promoters, chosen based on their predicted control via the transcriptional regulator NanR (*nanA, nanC, nanX and fimB*) using the Ecocyc database ^43^, were PCR amplified from *E. coli* NGF-1 gDNA using Q5 High-Fidelity PCR polymerase (NEB) following the manufacturer’s guidelines. PCR primers with type IIS RE sites (SapI) for individual sensors were designed using Primer 3 in Geneious Prime (software version 2020.1.1-2023.1.1; Biomatters Ltd). PCR products were purified using a Monarch PCR Cleanup Kit (NEB) before GGA assembly. Synthetic promoters EB1, EB2, EB3 and EB4 ^25^ were ordered as ssDNA and duplexed as dsDNA with SapI overhangs before GGA.

Reactions for GGA were set up with 3 nM of each DNA fragment component, 3 μL (15 units) of SapI, 0.25 μL (500 units) of T4 DNA Ligase, 2 μL of T4 DNA Ligase buffer and nuclease free water to a final volume of 20 μL. GGA reactions were cycled 30 times at 37°C for 5 minutes and 16°°C for 5 mins, then stored at 4°°C until transformation. 2 μL of the GGA mix was transformed into 50 μL of NEB 10-beta Competent E. coli (NEB) by heat shock according to the supplier’s protocol and plated onto carbenicillin and streptomycin selective LB-agar plates. After positive colony PCR, plasmids were miniprepped (Monarch® Plasmid Miniprep Kit, NEB) and transformed into electrocompetent E. coli NGF-1 encoding constitutive mKate2 fluorescence (DTR328). Transformant plates were scraped with pre-warmed LB, pooled, diluted 1:1000 and grown for 6 hours at 30°°C in LB-streptomycin-carbenicillin with 0.1% L-arabinose to induce Tn7-based genome integration. Cells were further back-diluted 1:1000 and grown overnight at 42°°C three times in order to cure the now genome integrated biosensors of the circuit plasmids. Individually cloned sensors were confirmed by sanger sequencing (Eurofins) using primers flanking the Tn7 insertion site ^42^.

### *In vitro* growth and induction of biosensor strains

Unless otherwise noted, bacterial strains were grown individually at 37°C in LB broth or on LB agar plates. Where appropriate, streptomycin and carbenicillin were supplemented at 50 μg/mL and 200 μg/mL respectively.

For *in vitro* sialic acid induction, bacterial cultures were diluted 1:500 (LB media) or 1:100 (M9 media + 0.4% glycerol). 100µL of diluted cultures were added to a 96-well plate with a row of either LB or M9 media, depending on the assay. The inducer, N-(-)-Acetylneuraminic acid, 97% (Neu5Ac) was added either at the time of dilution or after 5 hours of growth depending on the assay. Sensitivity test between Neu5Ac and other sialic acid variants was conducted using N-glycolylneuraminic acid (Neu5Gc), 2-keto-3-deoxynonulosonic acid (KDN), N-acetylmannosamine (manNAc) and N-Acetyl-2,3-dehydro-2-deoxyneuraminic acid (Neu5Ac2en; sialidase inhibitor) or the same amount of PBS in controls. Assays were performed in 96-well F-bottom black plates were covered with Breathe-Easy® sealing membrane (Sigma-Aldrich, Missouri, USA). On a CLARIOstar® Plus plate reader (BMG Labtech, Ortenberg, Germany), growing at 37°C, OD600 (optical density measured at a wavelength of 600nm) and fluorescence measurements (mVenus, Ex: 497-15/Em: 540-20; mKate2, Ex: 570-15/Em: 620-20) were taken every 15 mins with shaking between measurements. Fluorescence raw data was corrected for blank (media) by subtraction and normalised for OD600 and in some cases background fluorescence based on negative control strain DTR328 data to generate normalised fluorescence units. Fold change of normalised mVenus fluorescence was calculated based on the change of FU/OD600 at different concentrations of inducer within the same bacterial strain.

### Mouse models

All animal experiments were carried out under approval of the local Ethical Review Committee according to UK Home Office guidelines. 5-7 weeks old female C57BL/6 mice were purchased from Charles River (UK). Mice were provided with Teklad 2019 Global Rodent Diet (Envigo), alfalfa free, to reduce autofluorescence of gut contents during imaging. Upon arrival, mice were randomly distributed according to the group size of each experiment, followed by at least a week of acclimatation to the diet and the facility.

### *E. coli* NGF-1 engineered biosensor colonisation model

*E. coli* NGF-1 was administered as previously described ^27,28^. Mice were administered 5mg of streptomycin sulphate (VWR, 0382-EU-50G) in 100μL PBS by oral gavage. Plating of PBS diluted faecal samples on streptomycin selective LB agar plates 24 hours post streptomycin gavage confirmed the absence of background streptomycin resistant bacteria in the mouse gut. *E. coli* strains were grown overnight in LB, washed twice in PBS and concentrated 1:20 (∼10^11^ cells/mL). Mice were gavaged with 200 μL of the suspension. Bacterial load was estimated by serial dilution and plating of freshly collected faecal pellets that were resuspended to 100 mg/mL in PBS and vortexed for 2 minutes.

### Dextran sulfate sodium-induced colitis model

Mild-to-acute colitis was induced by oral administration of 1-3% DSS (36-50 kDa: MP Biomedicals) in the autoclaved drinking water of animals for 3 to 7 days depending on the experiment set up. To inhibit sialidase activity during or after DSS-induced colitis, 200 μL of N-Acetyl-2,3-dehydro-2-deoxyneuraminic acid (Neu5Ac2en) (10 mg kg-1 per day) was administered via oral gavage daily. Animals were monitored throughout the experiment for colitis severity through body weight, stool consistency and the presence of blood in faeces. At the end of the experiment, mice were humanely sacrificed, and the colons were retrieved for downstream analyses.

### Citrobacter rodentium infection model

Mice first received the engineered *E. coli* NGF-1 biosensor (DTR413) as previously described. Colonisation was confirmed by plating faecal samples on streptomycin-selective LB agar. After four days of colonisation, mice were challenged with ∼1 × 10⁹ CFU *Citrobacter rodentium* (ICC169) by oral gavage. Infection progression was monitored by daily weight recording and faecal CFU enumeration on day 1, 2, 4 and 6 post infection on nalidixic acid-selective LB agar for *C. rodentium* enumeration and streptomycin-selective LB agar for DTR413 CFU counts.

### Routine evaluation of inflammation

Retrieved whole colons were measured in length to evaluate the impact of inflammation. Hema-screen™ Single Slides (Avantor-VWR) were used to qualitatively detect occult bleeding due to inflammation where indicated. Scoring was performed blinded.

Faecal lipocalin-2 (LCN-2) levels were quantified as previously ^44^. Briefly, frozen faecal samples were reconstituted at 100 mg/mL in PBS containing 0.1% v/v Tween 20 and vortexed for 20 min to get a homogeneous suspension then centrifuged for 10 min at 12,000 rpm at 4°C. Clear supernatants were collected and stored at −20°C until analysis. LCN-2 levels were estimated in the supernatants by ELISA assay using the Duoset murine LCN-2 ELISA kit (R&D Systems) following the manufacturer’s protocols.

### Microbiota analysis

Faecal pellets (≈ 50 mg) were collected into 2ml microtubes with O-ring caps and stored at –80 °C. Prior to DNA extraction, samples were subjected to bead-beating by adding 100 mg of 0.1 mm zirconia beads and 1 ml of inhibiTEX buffer (Qiagen) to each tube. Samples were subjected to 3x 1 min bead-beating cycle using the Precellys Evolution homogenizer with 1 min rest on ice between. DNA extraction was then performed using QIAamp Fast DNA Stool Mini Kit as previously described ^45^.

For 16s rRNA sequencing, full length bacterial 16S rRNA genes (V1-V9) were amplified using PacBio forward (5’GCATC/barcode/AGRGTTYGATYMTGGCTCAG3’) and reverse (5’GCATC/barcode/RGYTACCTTGTTACGACTT3’) primers. DNA sample concentration was normalised at 500 pg/µL. KAPA HiFi HotStart 2x ReadyMix PCR was used for the reaction. Both the forward and reverse 16S primers were tailed with sample specific PacBio barcode sequences to allow for multiplexed sequencing. Barcoded 16S primers were arrayed into PCR plates using Echo acoustic liquid handler and the library was sequenced on Sequel IIe system on 1 SMRT cell 8M generating an average of 16,392 reads per sample (Q3: 20,765).

The PacBio sequencing data was processed using the nf-core/ampliseq pipeline, version 2.10.0 ^46^. Within this pipeline, data quality was evaluated with FastQC ^47^, summarized with MultiQC ^48^, and primers were trimmed using Cutadapt ^49^. Sequences were then examined as one pool (pseudo-pooled) using DADA2 to infer amplicon sequencing variants (ASVs). VSEARCH ^50^ was then used to cluster the ASVs with a pairwise identity of 0.97. Finally, taxonomic classification was performed by DADA2 and the database ‘Silva 138.1 prokaryotic SSU’ ^51^. Clustered ASVs and taxonomic identifications were then analysed using R with the vegan package ^52^ to preform permutational analyses of variance (PERMANOVA), beta dispersion analyses and nonmetric multidimensional scaling (NMDS) ordination. Plots were generated using ggplot ^53^.

### Gas Chromatography-Mass Spectrometry (GC-MS) of gut contents

Faecal pellets were extracted as previously described ^54^; all solvents were Optima HPLC Grade (Fisher) unless otherwise stated. Briefly, 400 µL chloroform was added to the pellet and vortexed, followed by 200µL methanol and vortexed again. Samples were centrifuged (4°C, 10 min, 16,000 rpm), the supernatant transferred to a clean Eppendorf 1.5 mL tube and dried using a rotary vacuum concentrator (Christ; 10 mbar, 30°C). 600 µL methanol:water (2:1 v/v containing 1 nmol of *scyllo*-inositol as an internal standard) was added to the sample pellet, centrifuged (4°C, 10 min, 16,000 rpm), and transferred to the previous dried extract. Chloroform (133 µL) and water (200 µL) were added to the sample, vortexed vigorously, and centrifuged as before. The upper, aqueous fraction was transferred to a glass vial and dried using a rotary vacuum concentrator as above. The dried sample was washed with 30 µL methanol and dried again, with this wash step repeated once more.

Polar metabolites were derivatised for GC-MS analysis by adding 20 µL freshly prepared methoxyamine hydrochloride (20 mg/mL in pyridine, both Sigma), followed by brief vortexing and incubation at room temperature overnight. Next, 20 µL BSTFA + 1% TCMS (‘TMS’) reagent (Sigma) was added to each sample and incubated for >1 hour before injection of 1 µL onto the GC-MS.

Data acquisition was performed largely as previously described ^55^, using an Agilent 7890B-7000C GC-triple quadrupole MS system, using the following parameters: carrier gas, helium; flow rate, 0.9 mL/min; column, DB-5MS (Agilent); inlet, 270°C; temperature gradient was 70°C (2 min), ramp to 295°C (12.5°C/min), and ramp to 320°C (25°C/min, 3 min hold). Scan range was m/z 40-550. Data was acquired using MassHunter software (version B.10.0.384.1). Data analysis was performed using MANIC software, an in house-developed adaptation of the GAVIN package ^56^. Qualitative analysis was performed using MassHunter Qualitative software (version 10.0). Metabolites were identified and quantified by comparison to authentic standards injected at the beginning and end of the sequence. Calibration curves for sialic acid (SA) linearity was calculated from 1, 10, 25, 50 and 100 µM standard solutions.

### Histological analysis of mouse colon

Swiss roll preparation of colon tissue was performed as previously described ^57^. Dissected colon tissue was placed on parafilm, surrounding fat tissue was removed, and the organ was open longitudinally. Gently, faecal pellets at distal colon were removed with forceps, preserving the mucus and digesta before the tissue was flattened with the luminal side facing up. The proximal extremity of the colon was rolled towards the distal end and a small needle (30G) was used to pierce all colon layers and keep the tissue rolled-up. Rolled tissues were immediately fixed at RT using PBS containing 4% PFA for 4h. After fixation, samples were washed three times in PBS and immediately placed in 15% sucrose in PBS (6-12h until the tissue sank) before being transferred to 30% sucrose in PBS overnight. The organs were place in a cryomold with OCT embedding matrix, snap-frozen in 2-methylbutane over dry ice and stored at –80°C until cryosectioning was performed.

Cryosectioning of OCT embedded tissue was performed using a Leica cryostat. 20 µm sections were cut and mounted on SuperFrost Ultra Plus™ GOLD slides. Samples for immunohistochemistry and *in situ* Hybdridisation were dried at RT for 10 min then fixed for 20 min in PBS containing 4% PFA follow by three washes to remove excess fixative.

Tissue for histological analysis was stained using hematoxylin (Mayer’s, Sigma, Gillingham, UK) and eosin (CellPath, Newtown, UK). Blind Histological analysis of tissue sections was then assessed and graded by a board-certified veterinary pathologist (ASB) of the Royal Veterinary College (London, UK) using light microscopy ^58,59^. Crypt architecture (0-4), inflammatory cell infiltrate (0-4), muscle thickening (0-4), goblet cell depletion (0-4) and crypt abscess (0-4) were independently scored and added together to give the overall histological score for three longitudinal regions (proximal, mid and distal colon) of each animal.

### Immunohistochemistry

For immunolabelling, the tissue was washed twice in PBS and incubated for 15 min in PBS containing 0.15% v/v Triton X-100 for permeabilization. Samples were then incubated in blocking solution 45 min (PBS containing 5% v/v donkey serum) followed by primary antibodies diluted in blocking solution (1:200) or *C. rodentium* anti-sera (1:50)^60^ for 1h at RT. Three washes in PBS ensued, before an incubation with secondary antibodies in blocking solution (1:500) for 45 min at RT. Triple immunofluorescence of host immune markers (CD68, CD45, CD3) was performed using the Opal multiplex staining system. Following each staining cycle, the Opal dye was covalently bound to the tissue, and antibody complexes were removed by heat-induced stripping, leaving the fluorescent Opal signal intact. For those slides where conjugated agglutinins or lectins were used, an extra incubation time of 1 hour after the secondary antibody was included. The percentage of area positive for CD68, CD45, CD3 and Ly-6G/Ly-6C was quantified by calculating the overlap between the automatically thresholded signal of each fluorescent channel and the DAPI-labelled regions. Analysis was performed separately within the proximal, mid, and distal regions of the swiss-roll colon.

After three washes to remove unbound antibody, PBS containing DAPI (1 µg/ml) was added for 10 min to label DNA. Finally, sections were washed three times in PBS and once with H2O before adding the mounting and closing the slides with coverslips.

### Microscopy analysis

*In vitro* activity of the sialic acid sensor, with and without antibodies, was evaluated in a Zeiss Axio Observer Widefield microscope with LED illumination, equipped with Plan Apo 100X/1.4 Oil Ph3 immersion objective, resolution 0,24 mm and controlled by Zen acquisition software and Hamamatsu Camera. The light Source was a Colibri 5/7, with LED 385 (32.1%) for DAPI, LED 475 (44.4%) for mVenus and LED 555 (60.5%) for mKate2 excitation.

Imaging of tissue samples was performed using a Leica SP8 inverted confocal microscope equipped with HCPL APO CS2 63X/1.40 Oil immersion objective (Resolution XY: 139.43; Resolution Z: 235.82; Free working distance: 140 mm), 4 Leica Lasers (Diode 405, Argon, DPSS 561, HeNe 594 and HeNe 633) and High sensitivity Detectors (HyD1, PMT2, HyD3 and PMT4).

Images were analysed using FIJI software. Quantification of reported levels was performed using a segmentation classifier generated in Labkit (FIJI) (Supplementary Figure 6) ^26^. The classifier was trained manually to distinguish between individual bacteria labelled for the constitutive fluorescence (mKate2) (all *in vivo* experiments) or based on DAPI labelled nuclei (*in vitro* experiments). Following a conservative approach, when clusters of bacteria could not be segmented individually, those particles were not included in the analysis. Segmented particles of between 0.5-2 mm^2^ were then used for average quantification of mVenus (*in vivo* experiments) or mVenus and mKate2 (*in vitro* experiments) signal. For illustration purpose, tile images were background subtracted 0.5 px, Gaussian blur 0.75 and Min/Max brightness adjusted in the same way for all samples within the experiment. Colour-blind friendly palettes were chosen based on the work of Okabe and Ito ^61^.

### Quantification and statistical analysis

Unless otherwise stated, statistical analyses were routinely performed by using GraphPad Prism (v10.0). Individual statistical tests along with correction for multiple comparisons used were chosen as appropriate to each dataset and are stated in the legend of each figure panel, accordingly.

## Supporting information

Supplementary Reagents and Resources Table

## Acknowledgements

We thank Eleanor Herbert and Alejandro Suarez-Bonnet for assistance with histopathology and Marko Storch for assistance with Pacbio 16s sequencing. The *E. coli* NGF-1 strain used was a kind gift from Pamela Silver. pSL1521 (pSPIN, pSC101* backbone) was a gift from Samuel H. Sternberg (Addgene plasmid # 160729; http://n2t.net/addgene:160729; RRID:Addgene_160729). The pRNA1-mKate2 template LPT36 was a kind gift of Laurent Potvin-Trottier and Johan Paulsson. Imaging was performed at The Facility for Imaging by Light Microscopy (FILM) at Imperial College London. The *C. rodentium* strain (ICC169) and anti-sera were a gift from Gad Frankel. This research was funded by a Sir Henry Dale Fellowship to D.T.R (211230/Z/18/Z). C.M.R. was funded by a President’s PhD Scholarship through Imperial College London.

## Author contributions

Conceptualization: D.C. and D.T.R.; Methodology: D.C., C.M.R., R.J., V. N., S.A.P-D., J.I.M., E.N. and D.T.R.; Investigation: D.C., C.M.R., P.L., V.N. and S.A.P-D; Formal Analysis: D.C., R.J., V.N., S.A.P-D, J.I.M. and D.T.R.; Writing – Original Draft and Visualisation: D.C. and D.T.R.; Writing - Review & Editing: D.C., C.M.R., P.L., J.I.M. and D.T.R; Supervision and Funding Acquisition: D.T.R.

