## Supplementary Figures 1-6 for "Bacterial whole-cell biosensors illuminate spatially variable sialic acid availability within the inflamed mammalian gut"

**[1] Department of Infectious Disease, Imperial College London, London, UK**

**[2] Francis Crick Institute, London, UK**

**[3] Imperial Centre for Engineering Biology, Imperial College London, London, UK**

**[4] Centre for Bacterial Resistance Biology, Imperial College London, London, UK**

[†] Current Address: A*Star Genome Institute of Singapore, Singapore

[‡] Current Address: Department of Women and Children's Health, King's College London, UK

[§] Current Address: Department of Food Science, Research Centre in Food and Development A.C., Hermosillo, Mexico

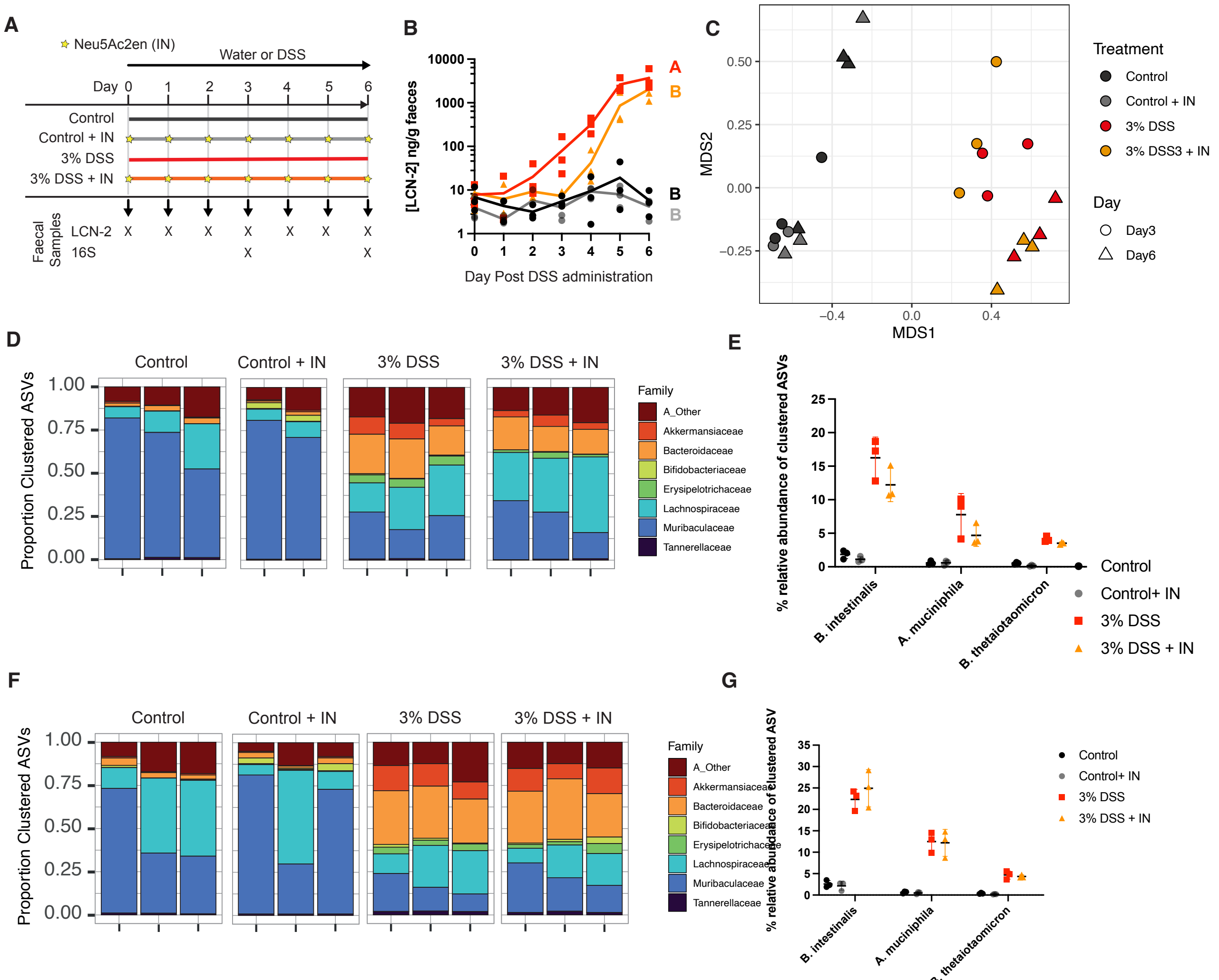


**Supplementary Figure 1.** A) To assess the impacts of inflammation and sialidase inhibition on the microbiome composition, DSS was administered in the drinking water of C57BL/6 mice (n=3 per group) at 0% (control) or 3% with or without daily oral gavage of Neu5Ac2en sialidase inhibitor (IN; stars). B) LCN-2 levels quantified by indirect ELISA of faecal pellets. Lines show mean. Statistics were calculated by one-way ANOVA with Tukey multiple comparison correction on AUC. Different compact letter (A-B) indicates significant differences between means. C) Long-read PacBio sequencing of the full 16s ribosomal RNA (rRNA) region followed by amplicon sequence variant (ASV) identification and clustering showed distinct microbiome compositions in DSS treated groups compared to controls (Pr(>F): 0.003). Our data were unable to distinguish between the microbiome in the presence or absence of sialidase inhibitor in either control (Pr (>F): 0.25) or DSS treatment groups (Pr(>F): 0.242). Multidimensional scaling of 16s rRNA sequencing data from faecal pellets at day 3 and 6 using monoMDS. Distance based on Bray Curtis, stress 0.0457 D) Differences were predominantly driven by bacterial families that have previously been associated with inflammation and DSS treatment, including increases in relative abundance of *Akkermansiacea* and *Bacteroidaceae* and decreases in *Muribaculaceae* Proportion of bacterial clustered ASVs within a family taxonomic classification at day 3. E) Percentage of relative abundance of clustered ASVs that showed similarity (over 97%) with those of *B. intestinalis*, *A. muciniphila* and *B. thetaiotaomicron* at day 3. F) Proportion of bacterial clustered ASVs within a family taxonomic classification at day 6. G) Percentage of relative abundance of clustered ASVs that showed similarity (over 97%) with those of *B. intestinalis*, *A. muciniphila* and *B. thetaiotaomicron* at day 6.

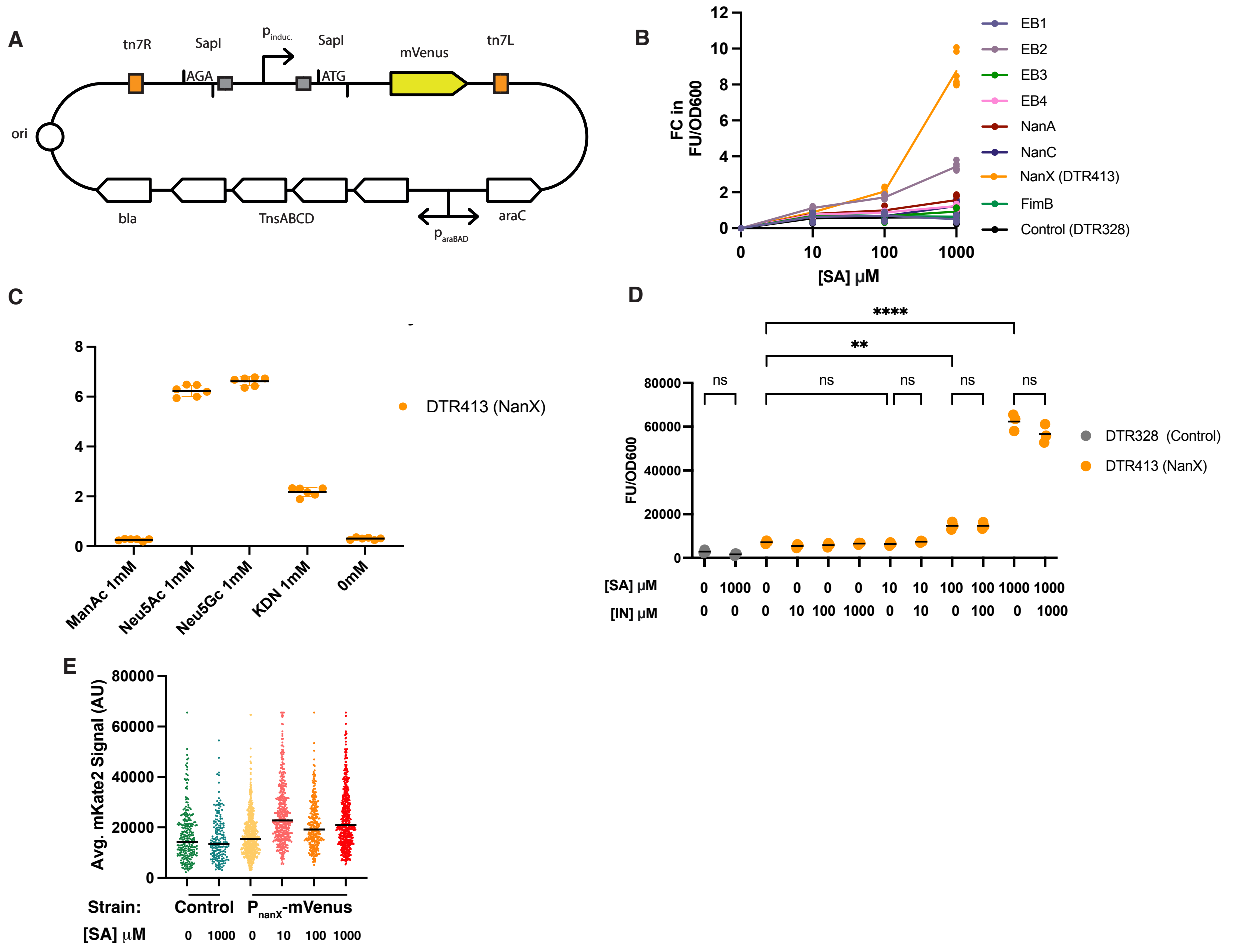


**Supplementary Figure 2.** A) Plasmid circuit design for genome integration of inducible promoters described in Figure 4. B) Fold change in fluorescence output normalised by OD600 at 6 hours exposed to different concentrations of Neu5Ac (SA; 10 µM, 100 µM or 1000 µM) in LB media compared to background fluorescence at 0µM. Signal for 8 putative sialic acid biosensors and a negative control (DTR328) lacking mVenus induction. Line shows mean of 3 replicates. C) Fold change in fluorescence output normalised by OD600 at 6 hours exposed to 1 mM of ManAc, Neu5Ac, Neu5GC, KDN or no supplementation (0 mM) in M9 media + 0.4% glycerol, compared to background fluorescence of negative control (DTR328). Line shows mean of six replicates. D) Fluorescence output of DTR413(NanX) and DTR328 (negative control), normalised by OD600, at 6 hours post induction in LB media supplemented with increasing concentrations of Neu5Ac and sialidase inhibitor Neu5Ac2en (IN). Graph shows mean of 3 replicates ± SD. Statistics shown for one-way ANOVA with Šídák multiple comparison adjustment. * = $p<0.05$, **= $p<0.01$, ***$p=<0.005$, ****$p=<0.0001$. E) Average intensity of mkate2 signal labelled with antibody (α-RFP) at increasing SA concentrations of 6h growth in the negative control (DTR328) and NanX (DTR413) evaluated by widefield microscopy as shown in Figure 2E-D.

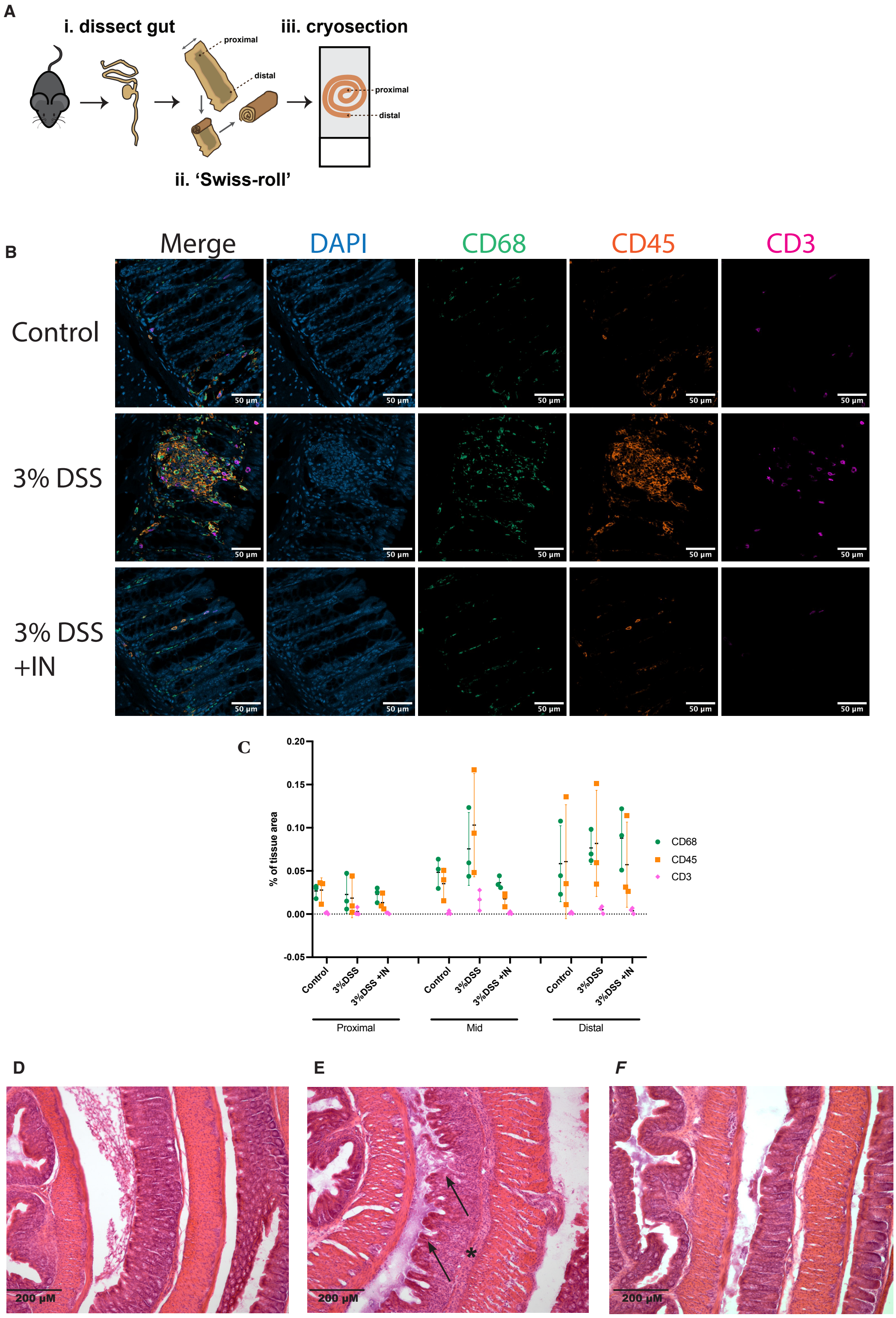


**Supplementary Figure 3**. A) Schematic of swiss-roll preparation of colon samples. B) Representative tile images of mid colon from Control, 3% DSS, and 3% DSS + IN treated animals of experiment described in Figure 5A. IHC labelling of monocyte-derived cells (α-CD68; green), Leukocytes (α-CD45, orange), and T-cells (α-CD3, magenta) along with nuclear staining (DAPI, blue) are shown. Scale bars are shown in figure. C) Percentage of area corresponding to cells positive for CD68, CD45 and CD3 in the three colon regions analysed individually. Line represents mean and SD bars. D-F) Representative H&E staining of Swiss-roll sections at the mid colon of Control (D), 3% DSS (E) and 3% DSS + IN (F) as used for histological scoring in Figure 3J. Arrows indicate Changes in crypt architecture and * highlight cellular infiltration in the submucosa. Scale bars are shown on the image.

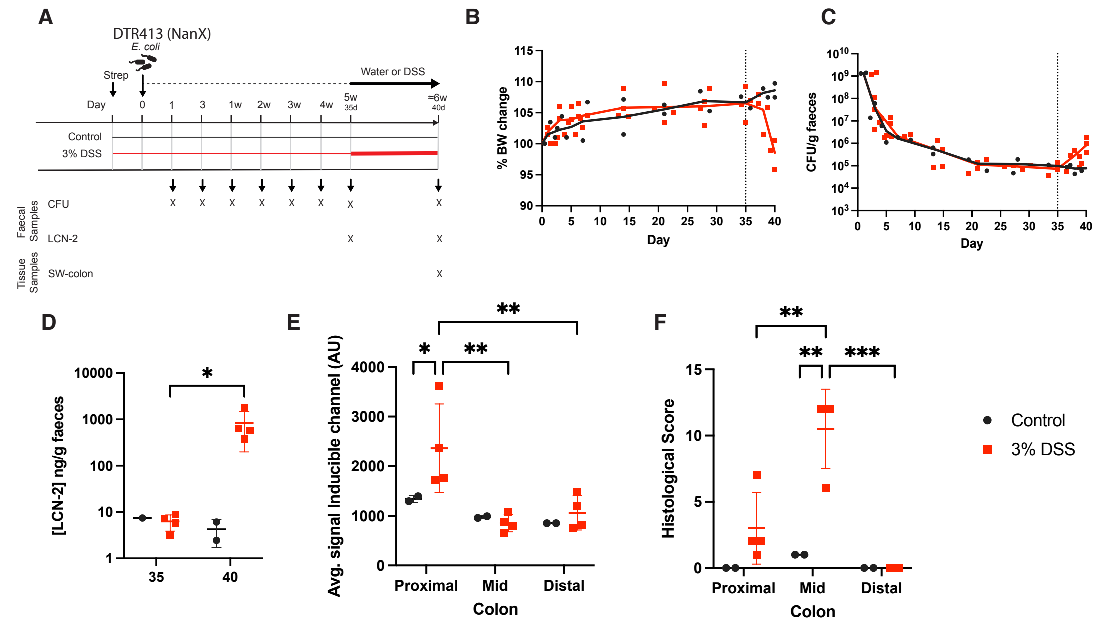


**Supplementary Figure 4.** A) DTR413 sialic acid-inducible whole cell biosensors were administered to C57BL/6 mice (n = 6) by gavage 24 hours following a single dose of streptomycin (5mg per mouse by oral gavage). After 35 days, 4 mice received DSS in the drinking water (3% for 5 days) while 2 mice remained as a control (0%). B) Body weight change, expressed as a percentage of starting body weight of each mouse. C) DTR413 abundance was measured in faecal pellets collected daily post administration. D) LCN-2 levels, as quantified by indirect ELISA in faecal pellets collected before (day 35) and after (day 40) DSS course. Graph shows mean ± SD. Statistics show results from two-way ANOVA uncorrected Fisher's LSD between samples on each day as shown. *=p<0.05. E) Geometric means of average signal of inducible mVenus reporter intensity (α-GFP) within bacteria segmented for constitutive (α-RFP) fluorochrome on swiss-roll sections on proximal, mid and distal colon whole colon at day 40. F) Histological Score in each gut regions from blinded analysis of H&E labelled sections, combining a sum of independently scored changes in crypt architecture, inflammatory infiltrate, muscle thickening, goblet cell depletion and crypt abscess. Lines show mean ± SD. For E-F, statistics show results from two-way ANOVA tests with Tukey multiple comparison adjustment comparing experimental groups within a gut region and between gut regions within each experimental group. between groups. * = p<0.05, **=p<0.005; ***p<0.0005.

A

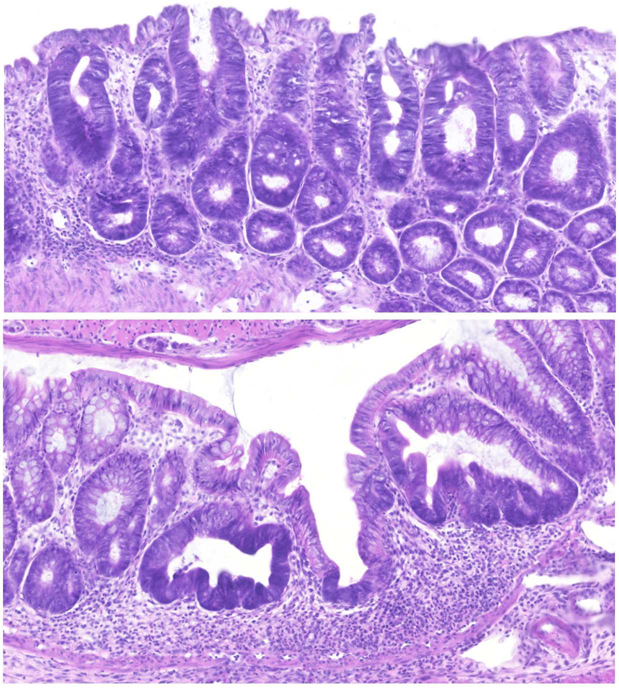

**Supplementary Figure 5.** A) Representative H&E staining of Swiss-roll colon samples of 3% DSS + IN of experiment described in Figure 5. Areas exhibit hyperplasia, increased cytoplasmic basophilia and increased mitotic activity, all features of regeneration after mucosal injury.

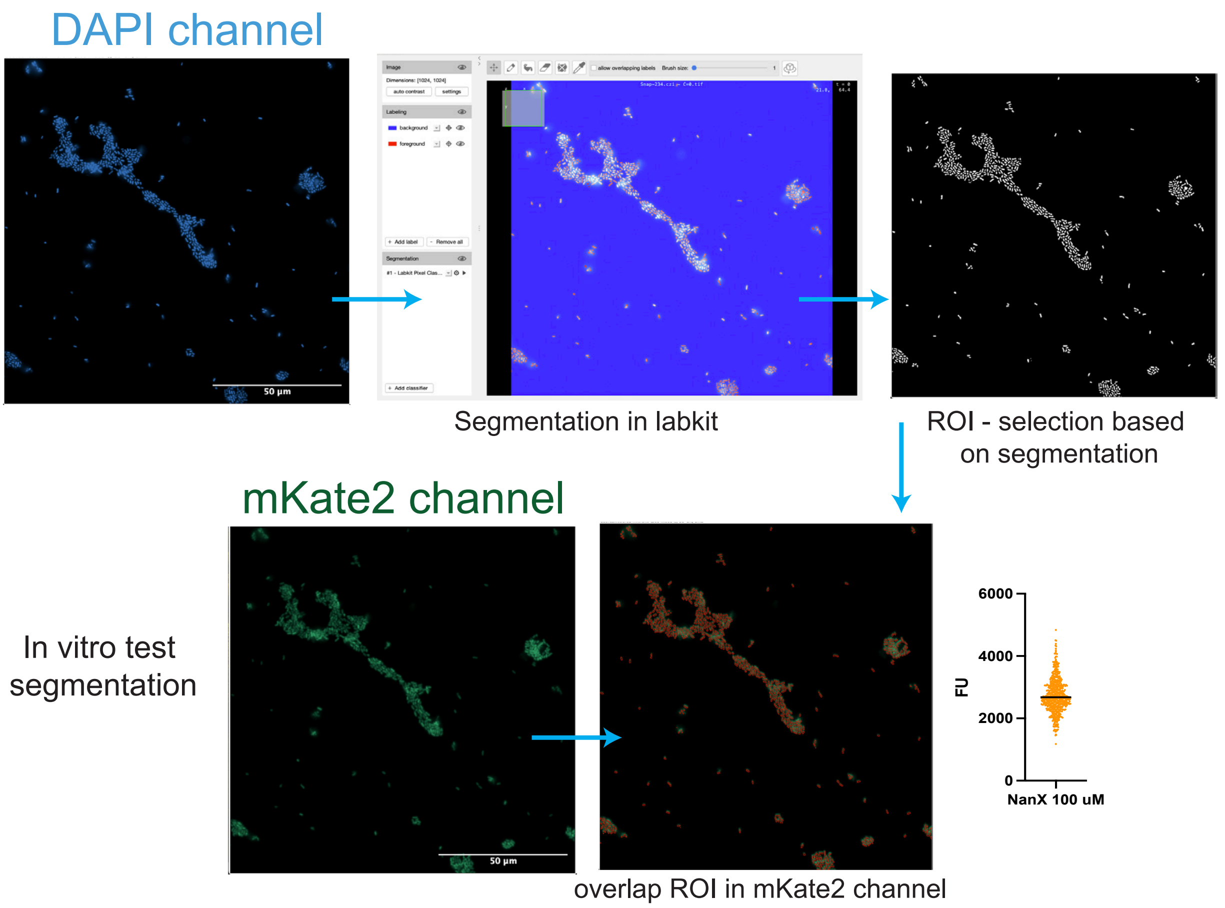


**Supplementary Figure 6** A) Workflow of segmentation analysis in Labkit software, FIJI. Training of classifier to create a selection ROI and overlay on the channel for average intensity analysis on the segmented areas.
